# Enhanced Ultrasound Image Formation with Computationally Efficient Cross-Angular Delay Multiply and Sum Beamforming

**DOI:** 10.1101/2024.09.03.611015

**Authors:** Cameron A. B. Smith, Matthieu Toulemonde, Marcelo Lerendegui, Kai Riemer, Dina Malounda, Peter D. Weinberg, Mikhail G. Shapiro, Meng-Xing Tang

## Abstract

Ultrasound imaging is a valuable clinical tool. It is commonly achieved using the delay and sum beamformer algorithm, which takes the signals received by an array of sensors and generates an image estimating the spatial distribution of the signal sources. This algorithm, while computationally efficient, has limited resolution and suffers from high side lobes. Nonlinear processing has proven to be an effective way to enhance the image quality produced by beamforming in a computationally efficient manner. In this work, we describe a new beamforming algorithm called Cross-Angular Delay Multiply and Sum, which takes advantage of nonlinear compounding to enhance contrast and resolution. This is then implemented with a mathematical reformulation to produce images with tighter point spread functions and enhanced contrast at a low computational cost. We tested this new algorithm over a range of *in vitro* and *in vivo* scenarios for both conventional B-Mode and amplitude modulation imaging, and for two types of ultrasound contrast agents, demonstrating its potential for clinical settings.

---

Receive Beamforming is a signal processing technique designed to estimate the spatial distribution of a series of sources. It is necessary to produce an image from the signals received by an array of sensors. In the field of medical ultrasound it is commonly achieved by utilizing the delay and sum (DAS) algorithm^1–3^, which attempts to solve this problem by calculating the time of flight from transmission to reception for a set of transducer elements to a specific location, finding the signal received by each element at this time point and then summing these signals before envelope detection. The intensity of the signal is then assigned to the pixel corresponding to that location before repeating the process for all pixels in a desired image. This algorithm, while computationally efficient, has limited resolution and suffers from high side lobes, limiting its effectiveness4.

One way to improve image quality is to take advantage of adaptive beamforming algorithms, many of which were originally developed for RADAR applications but have since been demonstrated to be effective for medical ultrasound. Most of these algorithms are based on the Capon beamformer^5–7^, which applies a data specific weighting to each element in the beamforming process in an attempt to minimise the contribution of noise and incoherent signals. While these algorithms improve image quality, they significantly increase computational cost.

A more recent set of beamformers use nonlinear processing to improve image quality in a more computationally efficient manner^4,8–11^. A major contribution was the Delay Multiply And Sum (DMAS) algorithm^4^, which cross-multiplies the signals received by different elements by combinatorically coupling elements. This can be interpreted as the aperture auto-correlation function, computing the spatial cross-correlation among the received signals. In doing so, the DMAS algorithm successfully reduces the side lobes and improves image quality.

Later, the Frame Multiply And Sum (FMAS) algorithm^8^ was developed and applied to ultrafast power Doppler imaging. This algorithm utilises nonlinear angular compounding. It performs conventional DAS on each angle transmission in plane-wave ultrafast imaging and then cross-multiplies pairs of acquisitions to evaluate the spatial coherence in the generated radio frequency (RF) images. This improves the contrast-to-noise ratio over DAS at a computational cost only marginally higher than DAS as the number of angles is usually small.

In this work, we present a new algorithm called Cross-Angular Delay Multiply and Sum (CADMAS), which improves the image quality of ultrafast imaging by evaluating the coherence between all received signals across all acquisitions at different angles. We additionally take advantage of a mathematical reformulation, decreasing the computation time compared to conventional implementations of DMAS, to create a beamformer that enhances image quality at a low computational cost.

The CADMAS algorithm can be defined by:

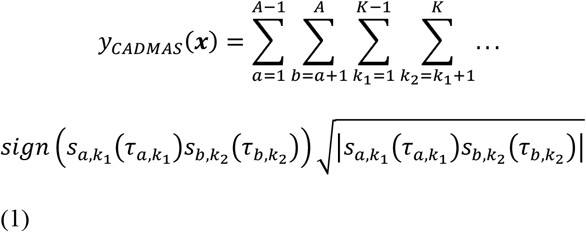

where A is the total number of angles, K is the total number of transducer elements, s is the signal received at a specific element and transmitted angle τ, is the time delay accounting for the time of flight for a specific transmission angle and element to position τ, and is the pixel intensity allocated to position τ. The schematic for this algorithm can be found in figure 1. In this form, the CADMAS algorithm, while easily parallelizable, contains a number of iterations based on the number of cross-angle-element pairings, which can lead to a high computational cost. However, by taking advantage of a mathematical reformulation similar to that described by Polichetti *et al*.^11^, the algorithm can be simplified computationally utilizing:

**FIG. 1.**
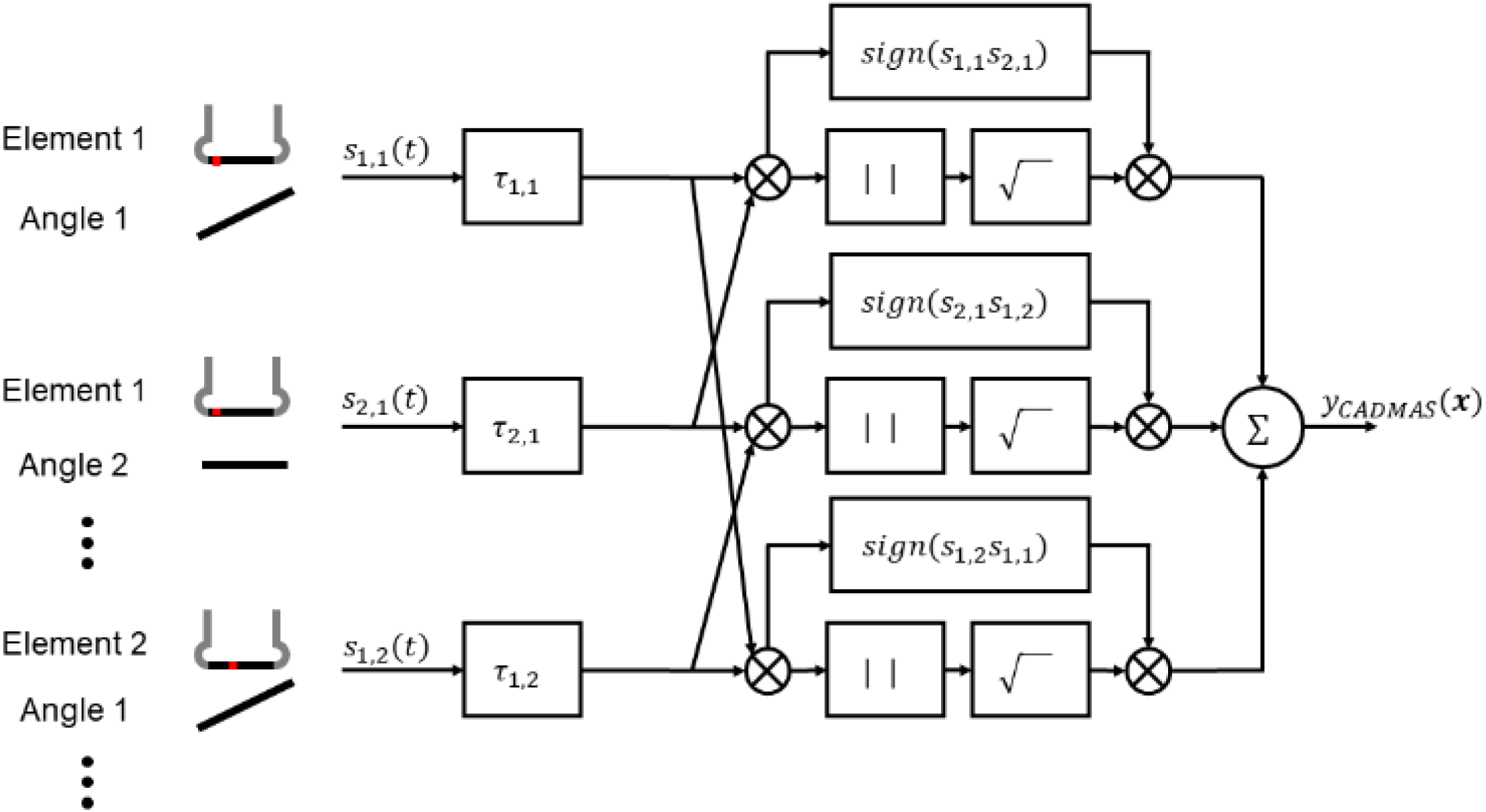
Block diagram showing how the CADMAS algorithm combines recorded signals from different elements and angle transmissions to produce the desired image.

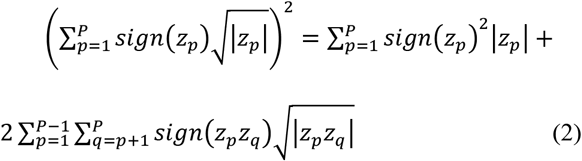

where is an element of any vector of length.

Therefore

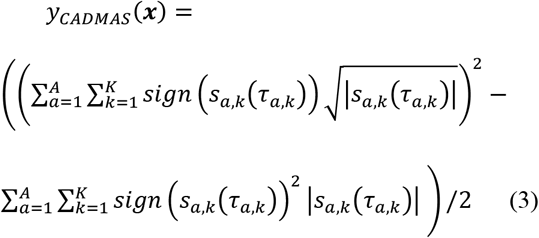

This formulation brings the computation time down to 1-2 fold that of DAS, depending on the implementation and hardware, with a similar computation time to DMAS if the latter is implemented utilising the same reformulation.

To investigate the effectiveness of the CADMAS algorithm, we conducted a series of experiments in which CADMAS was compared to DAS, FMAS, and DMAS, in addition to DMAS+FMAS, in which DMAS beamforming is implemented for each angle and then angles are compounded with the FMAS algorithm. Initially, we studied the effect of CADMAS on the point spread function of a wire scatterer. We imaged a 50 μm wire at a distance of 26 mm from a GE L3-12-D linear array transducer operated with a Vantage 256 Verasonics scanner with a transmitted frequency of 5 MHz, 0.1 MI, 0.5 cycle. We found that the point spread function of CADMAS had significantly reduced side-lobes and noise floor compared to the other beamforming schemes (Fig. 2).

**FIG. 2.**
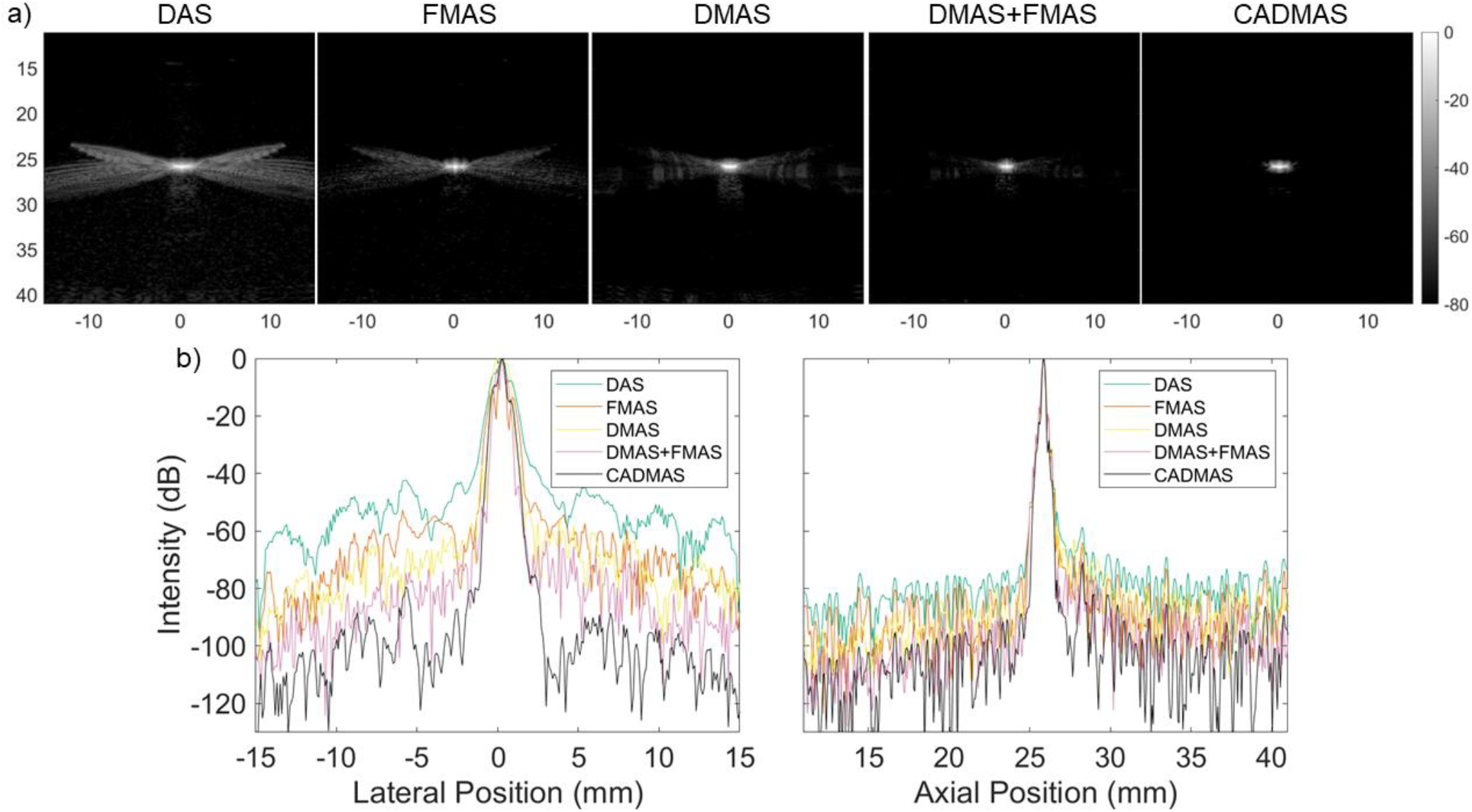
(a) Images obtained by beamforming an experimental wire scatter dataset in five different ways. (b) Lateral and axial cross sections of the images for comparison.

To quantify the improvement in image contrast in a controlled in vitro environment, we utilized a GAMMEX phantom (GAMMEX Ultrasound 403 LE multipurpose phantom) using a GE L3-12-D array with a transmitted frequency of 5 MHz, 0.3 MI, 1.5 cycles. The images produced by the five algorithms can be seen in Fig. 3a. We calculated Contrast-to-Tissue Ratios (CTRs) to allow for quantitative comparison, according to:

**FIG. 3.**
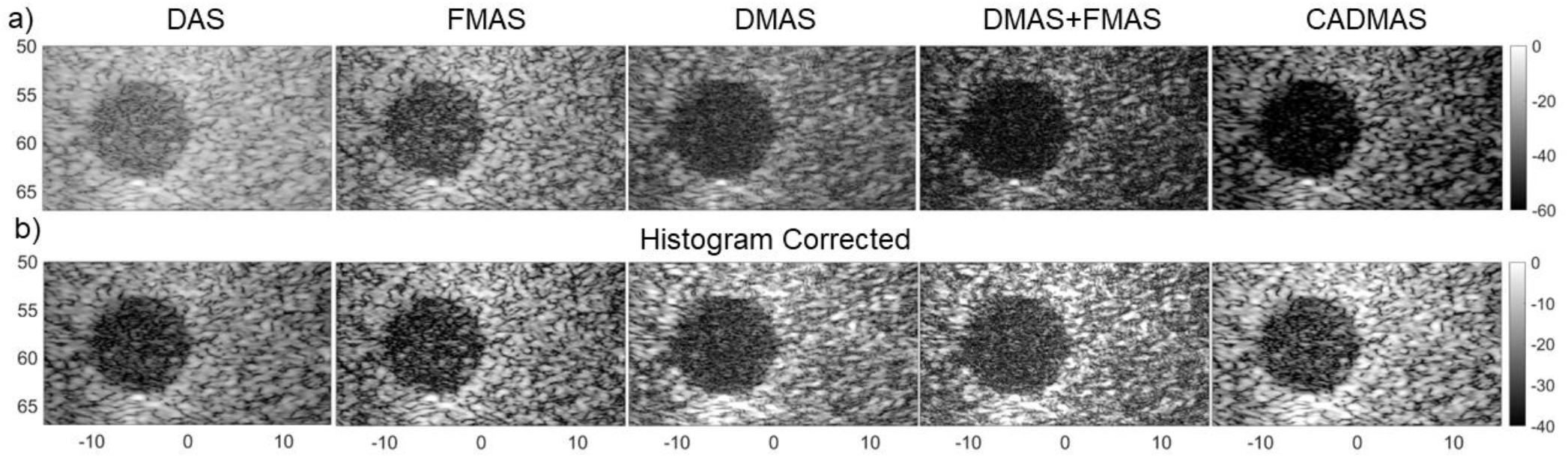
(a) Images obtained by beamforming a GAMEX phantom void dataset in five different ways. (b) The same images after histogram correction.

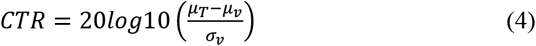

where is the mean of the pixel intensities in the tissue, is the mean of the pixel intensities in the void and is the standard deviation of the pixel intensities in the void.

Quantitative comparison of images with different beamforming methods is difficult to conduct fairly. Many metrics and methods have been derived, each having strengths and weaknesses^12–14^. A key issue is the effect of nonlinear shifts in the dynamic range, which can artefactually affect metrics such as the commonly used CTR. In an attempt to conduct a fair comparison, we calculated the CTR for both our original images (Fig. 3a) and for images generated using the partial histogram matching technique described by Bottenus *et al*^14^ (Fig. 3b). CTR values were higher for CADMAS compared to other beamformers regardless of scaling (Table I).

**TABLE 1.** Mean and standard deviation of contrast-to-noise ratio when beamforming GAMMEX void phantom datasets in five different ways (n = 10)

|  | *DAS* | *FMAS* | *DMAS* | *DMAS+FMAS* | *CADMAS* |
| --- | --- | --- | --- | --- | --- |
| CTR | 16.0 ± 0.2 | 16.2 ± 0.4 | 18.8 ± 0.5 | 16.8 ± 0.6 | 23.6 ± 0.7 |
| CTR (Histogram Corrected) | 16.0 ± 0.2 | 14.9 ± 0.4 | 18.7 ± 0.5 | 15.5 ± 0.5 | 19.4 ± 0.5 |

To test the CADMAS algorithm for use with amplitude modulation (AM), a contrast-specific mode initially designed to image nonlinear scatters, a 2 mm diameter cylinder of gas vesicles (GVs) set inside a tissue mimicking phantom comprising 1% (w/v) agar (Agar W201201, SigmaAldrich), 0.2% (w/v) Al_2_O_3_ (White Aluminium Oxide Powder Fepa Grit: 1200, 3 Micron, Kemet) was imaged using an L11-4v (Verasonics) with a transmitted frequency of 4 MHz, 1 cycle, pressure ranging from 120 to 540 kPa peak positive pressure. GVs are protein-shelled, air-filled sub-micron particles^15^ naturally expressed by buoyant photosynthetic microbes_16_. They can be purified as nanoscale contrast agents or expressed heterologously in bacterial and mammalian cells as acoustic reporter genes^17–19^. GVs provide strong contrast to tissue using AM imaging^20,21^. We acquired images of GVs purified from *Anabaena flos-aquae*^22,23^ (Fig. 4a shows a representative image at 420 kPa) and analysed the CTR at each transmit pressure according to:

**FIG. 4.**
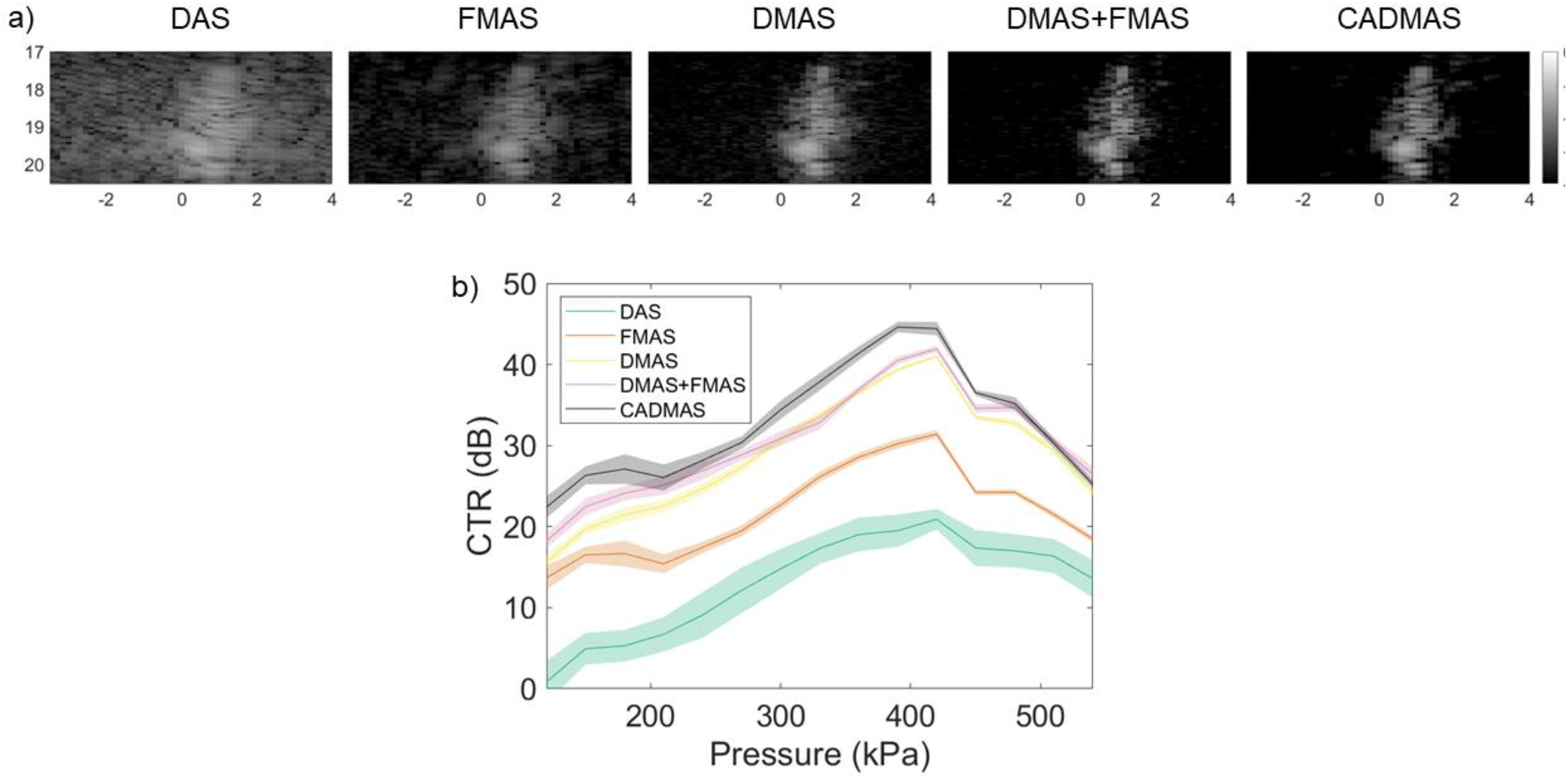
(a) Images obtained by beamforming an AM dataset imaging gas vesicles embedded in a tissue phantom at 430 kPa peak negative pressure in five different ways. (b) Mean and standard deviation of the CTR calculated for each beamforming technique throughout the applied pressure ramp (n=10).

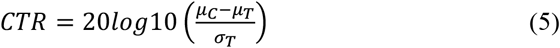

where is the mean of the pixel intensities in the tissue, is the mean of the pixel intensities in the void, and is the standard deviation of the pixel intensities in the tissue over a series of acquisitions. As expected from the nonlinear mechanical behaviour of GVs, CTR increased with pressure until a pressure was reached at which the GVs started to collapse irreversibly and the CTR began to decrease _24_.

The ability of CADMAS to improve *in vivo* images was investigated by analysing images of a rabbit kidney. The experiment complied with the Animals (Scientific Procedures) Act 1986 and received approval from the Animal Welfare and Ethical Review Body of Imperial College London. A pathogen-free male New Zealand White rabbit (HSDIF strain, Envigo, Huntingdon, UK) aged 20 weeks and weighing 3.14 kg was maintained on a 12:12 h light:dark cycle at 18°C and fed a standard laboratory diet. It was pre-medicated with acepromazine (0.5 mg/kg, i.m., CVet, London, UK), anaesthetised using a combination of medetomidine (Domitor, 0.25 mL/kg, i.m.) and ketamine (Narketan, 0.15 mL/kg, i.m.) and maintained with a third of the initial dose of anaesthetics every 30 to 45 minutes for up to 4 hours. The rabbit underwent tracheotomy and was ventilated at 40 breaths/minute. It was positioned supine on a heated mat with, continuous monitoring of heart rate and blood oxygen saturation, and fur was removed from the abdominal region. The left kidney was imaged transcutaneously during after the injection of microbubble contrast agents via the marginal ear vein; an initial dose of 0.1 mL was followed by a second of 0.05 ml. The perfluorobutane microbbubles were prepared in-house by dissolving 1,2-Dipalmitoylsn-glycero-3-phosphatidylcholine, 1,2-dipalmitoyl-sn-glycero-3-phosphatidyleth-anolamine-polyethylene glycol 2000 and 1,2-dipalmitoyl-3-trimethylammonium propane in water with a molar ratio of 65:5:30 and a total lipid concentration of 0.75, 1.5 and 3 mg/mL respectively. This resulted in a solution with 15% propylene, 5% glycerol and 80% saline. Perfluorobutane was injected into a 2 ml container containing 1.5 ml of the prepared solution just before use and the microbubbles, were activated by agitation in a shaker for 60 seconds, giving a final concentration of 5 × 10^9^microbubbles/mL^25^. Euthanasia employed pentobarbital (0.8 mL/kg). We imaged the kidney transcutaneously using a GEL3-12-D probe with a transmitted frequency of 5 MHz, 0.1 MI, 1 cycle, 100 frames per second, 10 angles ±10°.

In images accumulated from 500 sequential frames (Fig. 5), the CADMAS beamformer enhanced contrast while suppressing incoherent signals and yielded the highest CTR across the five imaging methods (Table II). The method could also be applied to each individual frame, allowing easier microbubble segregation, and hence improved super-resolution images, although that was outside the scope of this work.

**FIG. 5.**
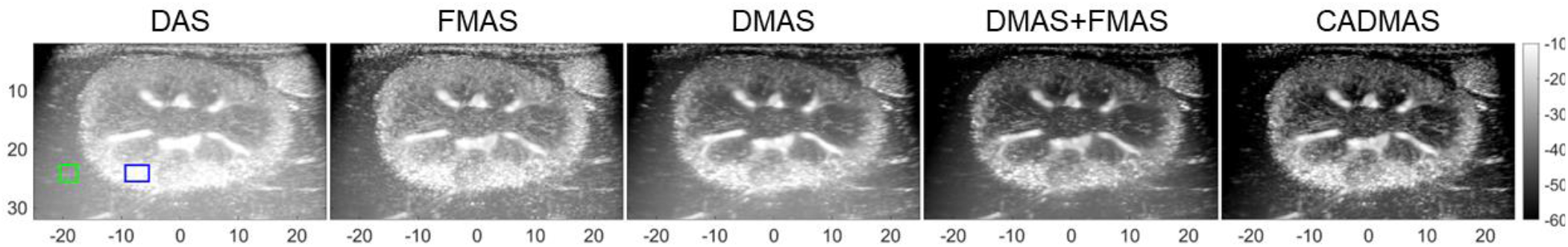
Images accumulated over 500 frames by beamforming a rabbit kidney dataset in five different ways, Contrast and tissue regions used for CTR estimations are designated by blue and green boxes, respectively.

**TABLE 2.** Contrast-to-noise ratio when beamforming an in viv*o* rabbit kidney dataset in five different ways.

|  | *DAS* | *FMAS* | *DMAS* | *DMAS+FMAS* | *CADMAS* |
| --- | --- | --- | --- | --- | --- |
| CTR | 33.9 | 37.7 | 39.8 | 44.3 | 46.1 |

In summary, the CADMAS algorithm provided a tighter point spread function and greater contrast than DAS, DMAS, FMAS, and DMAS+FMAS, and at low computational cost. The improvements were demonstrated in both conventional B-Mode and AM imaging, with GV and microbubble contrast agents, and for *in vitro* and *in vivo* imaging. The results suggest that CADMAS has potential utility in biological research and clinical settings.

## ACKNOWLEDGMENTS

The authors gratefully acknowledge funding from the International Human Frontier Science Program Organization (grant LT0036/2022-L), the Engineering and Physical Sciences Research Council (Grant No. EP/T008970/1 and EP/V04799X/1), the National Institute for Health Research i4i (Grant NIHR200972), the National Institutes of Health (R01NS120828 to M.G.S.), the Chan Zuckerberg Initiative, and the British Heart Foundation Centre for Research Excellence (grant PG/16/95/32350).

